# Functional Equivalence of Heat-Inactivated (HI) and Live Probiotic RSB11 in Suppressing Inflammation: Expanding Formulation and Application Potential

**DOI:** 10.64898/2026.02.11.705228

**Authors:** Teodora Nicola, Tanmaya Madhvacharyula, Ajay Ashok, Aryan Mandot, Iman Abdelgawad, Ryann Singh, Kara Siedman, Youfeng Yang, Namasivayam Ambalavanan, Charitharth Vivek Lal

**Affiliations:** Division of Neonatology, Department of Pediatrics, University of Alabama at Birmingham, Birmingham, AL, USA; Department of Ophthalmology, Schepens Eye Research Institute of Massachusetts Eye and Ear, Harvard Medical School, Boston, MA 02114, USA; Lung Health Center, University of Alabama at Birmingham, Birmingham, AL, USA; Marnix Heersink Institute of Biomedical Innovation, University of Alabama at Birmingham, Birmingham, AL, USA

## Abstract

The clinical potential of probiotics has been widely recognized, but their translation into reliable therapeutic products has been hindered by major limitations such as undesirable immunogenic responses, the need to maintain viability, instability during storage and transport, and concerns regarding safety in vulnerable populations. Postbiotics, defined as inanimate microbial cells or their components with pro-health activities, overcome many of these limitations by offering enhanced stability, reproducibility, and safety. However, it is very vital to understand if the heat inactivation (conversion of a probiotic to its postbiotic inert form) compromise its functional efficacy. Here, we systematically compared a novel probiotic-derived candidate, *Lactiplantibacillus plantarum* RSB11 strain, in its live (RSB11 Life, probiotic) and heat-inactivated (RSB11-HI, postbiotic) forms across multiple human epithelial and non-epithelial models relevant to inflammation driven pathologies. To investigate the gut-tissue(s)-axis concept we used gut (Caco-2), lung (HBE), ovary (BG1), bone (osteoblasts, MG-63), kidney (A-498) and liver (HepG2) cells exposed to *E-coli* or lipopolysaccharide (LPS), and quantified matrix metalloproteinase-9 (MMP-9), an inflammatory mediator, by qPCR and pro-inflammatory cytokines such as tumor necrosis factor-α (TNF-α), IL-6, and IL-1β by ELISA. In addition, we assessed β-glucuronidase activity and estrogen modulation to explore gut–ovarian axis signaling. Across all models, both RSB11 Life and RSB11-HI robustly suppressed MMP-9, TNF-α, IL-6 and IL-1β induction, with equivalent magnitude of effect. The inactivated form retained full cytokine-suppressive capacity and, notably, enhanced β-glucuronidase activity, suggesting additional benefits in microbiome hormone cross-talk. Our findings demonstrate that heat inactivation does not compromise, and may even expand, the functional range of RSB11. By maintaining bioactivity while eliminating the drawbacks of live biotics, heat inactivated RSB11 emerges as a robust, scalable, and versatile postbiotic with potential applications in systemic inflammatory disorders.

**Graphical Abstract:** 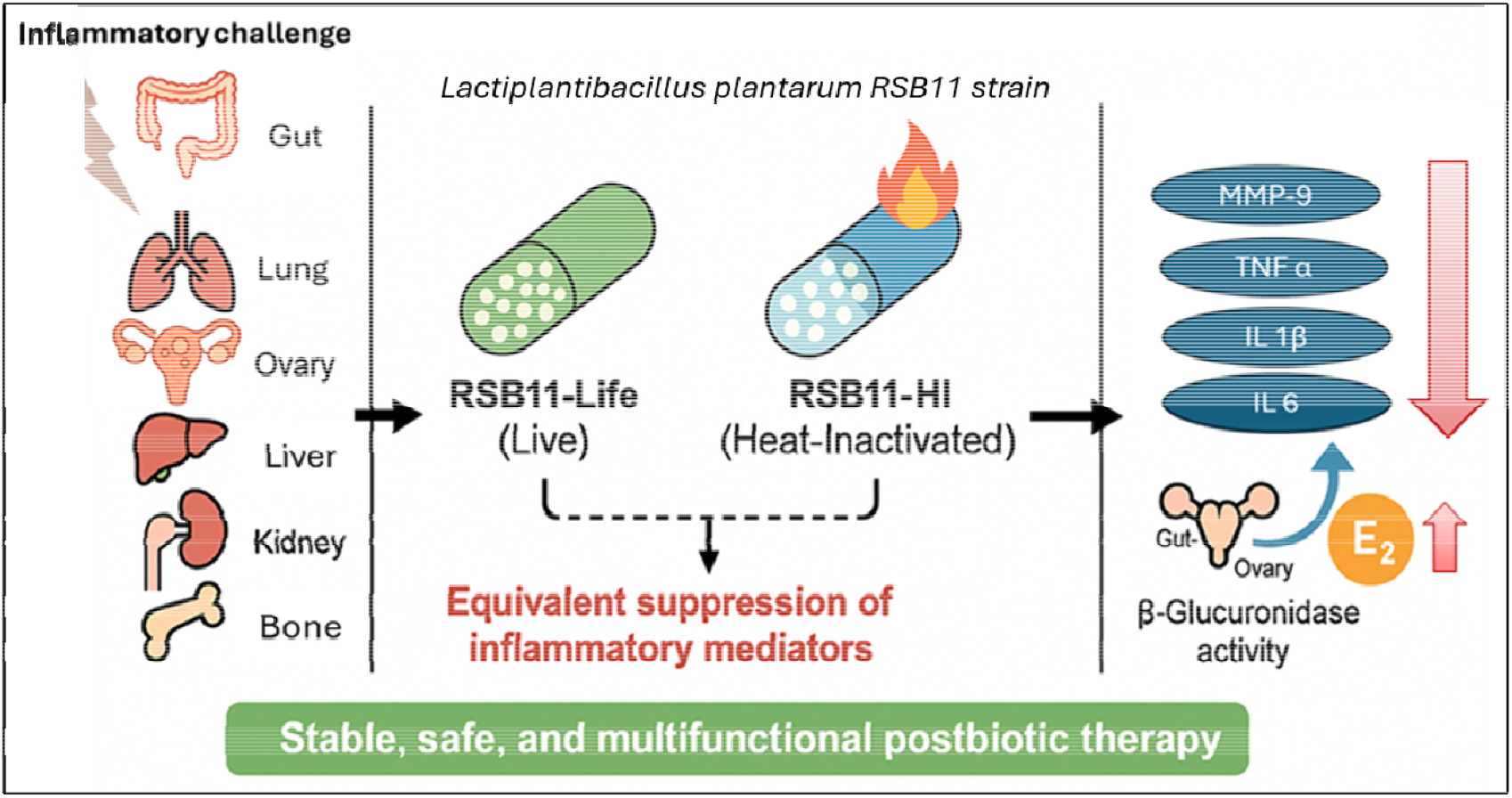

**Graphical abstract of RSB11-HI activity:** Heat-inactivated postbiotic RSB11-HI retains the anti-inflammatory efficacy of its live counterpart (RSB11Life) across diverse organ-relevant cell models. Upon LPS or *E-coli* stimulation, epithelial, immune, and tissue-specific cells (gut, lung, ovary, bone, kidney, liver) upregulate pro-inflammatory mediators including MMP-9, TNF-α, IL-6, and IL-1β. Both RSB11Life and RSB11-HI effectively suppress these inflammatory responses, with RSB11-HI exhibiting more consistent and robust reductions of inflammatory markers across models. Additionally, RSB11-HI uniquely enhances β-glucuronidase activity, facilitating estrogen metabolism and signaling through the gut–ovary axis. Together, these findings highlight RSB11-HI as a stable, safe, and multifunctional postbiotic candidate suitable for therapeutic formulation. *Image was designed using ChatGPT*.

## Introduction

Chronic, low-grade inflammation underlies the pathogenesis of a wide spectrum of human diseases, including gastrointestinal disorders, chronic respiratory disease, metabolic syndromes, reproductive dysfunction, liver fibrosis, kidney disease, and neurodegeneration. Central to these processes are inflammatory mediators such as tumor necrosis factor-α (TNF-α)[1], interleukin-6 (IL-6)[2], interleukin-1β (IL-1β)[3] and matrix metalloproteinase-9 (MMP-9)[4], which amplify cytokine cascades, promote extracellular matrix remodeling, and disrupt tissue homeostasis[5]. While pharmacological agents targeting these pathways exist, they are often costly, associated with systemic side effects, or limited to specific disease indications. This has spurred growing interest in microbiome-based interventions as safe, broadly acting strategies for dampening inflammation and restoring host–microbiome balance[6; 7; 8].

Probiotics have been extensively studied in this context and have shown immunomodulatory effects in both preclinical and clinical settings[7; 9; 10; 11]. However, their widespread adoption as therapeutics has been hindered by major practical and biological constraints[12; 13]. Chief among these is the reliance on viability since live microorganisms require precise storage conditions[14], are vulnerable to degradation when mixed with supplemental compounds[15], and exhibit unpredictable survival within the gastrointestinal tract. These issues surrounding the use of live forms of microorganisms bring variability in efficacy across batches and formulations, undermining reproducibility and reliability. Moreover, safety concerns have been raised regarding the administration of live bacteria in immunocompromised, critically ill, or neonatal populations, where translocation or overgrowth could pose risks[16; 17; 18]. Thus, while probiotics remain conceptually attractive, the requirement for microbial viability limits their translational feasibility and scalability.

Postbiotics, defined as preparations of inanimate microorganisms and/or their bioactive components that confer health benefits to the host have emerged as a solution to these limitations[19; 20]. By decoupling efficacy from viability, postbiotics offer enhanced stability, reproducibility, and safety. They can be stored without refrigeration, formulated into diverse delivery systems, and used in populations where live organisms are contraindicated[21]. Despite this promise, a key unresolved question has been whether heat inactivation compromises function and efficacy of these formulations. Exploring the efficacy of heat-inactivated (HI) formulations is therefore critical. If shown to be functionally equivalent to live probiotics, HI preparations would overcome barriers that currently restrict probiotic application. They would combine therapeutic potency with superior stability and safety, unlocking avenues for incorporation into capsules, powders, beverages, and even topical applications without fear of potency loss. Demonstrating equivalence between live and HI forms thus represents not just an incremental advancement but a shift in standards in microbiome-based therapy.

In the present study, we address this critical gap by comparing the effects of a candidate postbiotic, RSB11, in both its live (RSB11-Life) and heat-inactivated (RSB11-HI) forms[11; 22]. Using a panel of human epithelial and non-epithelial cell models representing the gut[4; 23], lung[24], ovary[25], bone[26], kidney[27] and liver[28], we examined their ability to suppress MMP-9, TNF-α, IL-6 and IL1β expression in response to inflammatory challenge with *E-coli* or lipopolysaccharide (LPS). We further assessed metabolic cross-talk by evaluating β-glucuronidase activity and estrogen modulation along the gut–ovarian axis. By deconjugating glucuronidated estrogens, β-glucuronidase facilitates their reactivation and recirculation, thereby influencing local and systemic estrogen bioavailability[29]. Assessing β-glucuronidase activity provides functional insight into how RSB11-HI may restore microbial endocrine balance and modulate estrogen-dependent inflammatory responses in ovarian and gut epithelial systems. By demonstrating that heat-inactivated RSB11 retains full immunomodulatory efficacy, this work establishes a foundation for the development of RSB11-HI as a next-generation postbiotic therapeutic for inflammation-driven disorders.

## Materials and Methods

### Reagents and Materials

Lipopolysaccharide (LPS)[30] and *E-coli* (10^7^ CFU/ml)[31] were used as inflammatory stimuli. Unless specified, molecular biology reagents (RNeasy Mini Kit, iScript™ cDNA kit, TaqMan master mix, all from Thermo Fisher) and ELISA kits (R&D Systems) were from standard vendors. Microplates were read on a BioTek Epoch reader.

### Cell Lines and Culture Conditions

Unless specified, all cell lines were purchased from American Type Culture Collection (ATCC). Human lung bronchial epithelial cells (16HBE, Sigma) were cultured as per manufacturer’s recommendations at 37 °C, 5% CO_2_, with medium changes every 2–3 days. Caco-2 (human colorectal adenocarcinoma) were maintained in DMEM with 10% FBS, 1% penicillin–streptomycin, and 1% non-essential amino acids at 37 °C, 5% CO_2_. MG-63 (human osteoblast-like) and (BG1) ovarian cells were cultured in EMEM with 10% FBS (Thermo Fisher) and 1% antibiotic–antimycotic (Thermo Fisher). HepG2 (human hepatocellular carcinoma) were maintained in MEM (Thermo Fisher) with 10% heat-inactivated FBS, 1% non-essential amino acids, and 1% penicillin– streptomycin. A-498 (human renal carcinoma) were grown in RPMI-1640 (Thermo Fisher) with 10% FBS and 1% antibiotic–antimycotic. Cells were sub-cultured at ∼70–90% confluence according to each line’s standard practice.

**Table 1.** Cell lines and study design.

| Organs | Cell lines | qPCR | ELISA |
| --- | --- | --- | --- |
| Gut | Caco-2 | MMP9 | TNFα, IL-6, IL-1β |
| Lung | HBE | MMP9 | TNFα, IL-6, IL-1β |
| Ovary | BG1 | MMP9 | TNFα, IL-6, IL-1β |
| Bone | MG63 | MMP9 | TNFα, IL-6, IL-1β |
| Liver | HepG2 | MMP9 | TNFα, IL-6, IL-1β |
| Kidney | A498 | MMP9 | TNFα, IL-6, IL-1β |

### Availability of the Heat-Inactivated Postbiotic (RSB11-HI)

The life and heat-inactivated *Lactiplantibacillus plantarum* RSB11 strain was kindly provided by Resbiotic Nutrition Inc. (resbiotic.com)[32]. Several concentrations were tested, and a working concentration of 3 mg/mL was sufficient to produce the desired experimental outcome. The freeze-dried powders, both live and heat-inactivated, were stored at -80°C to ensure the long-term stability of the biological material. In this study, the term “heat-inactivated” refers to a heat-treated preparation produced under controlled thermal conditions to ensure safety and functional stability, with process validation currently being finalized by the manufacturer.

### Inflammatory Stimulation and Treatment Paradigms

Cells were seeded in 24-well plates and grown to ∼80% confluence. Inflammation was induced with LPS (5 µg/mL) or by *E-coli* (10^7^ CFU/mL) incubation for 4 h. For each cell line, the following groups were run in parallel: untreated control; LPS/*E-coli* only; RSB11-life only, RSB11-HI only; LPS/*E-coli* + RSB11-life and LPS/*E-coli* + RSB11-HI. For the gut-ovarian axis the supernatants from previously treated Caco-2 cells were collected and used for 4hrs treatment of BG1 ovarian cells while equal volume of fresh EMEM media was added.

### RNA Isolation and Quantitative PCR

Total RNA was isolated (RNeasy Mini Kit, Qiagen) and reverse-transcribed (iScript™, Bio-Rad). qPCR was performed using TaqMan Master Mix (Thermo Fisher Scientific) on a QuantStudio 3 system per manufacturer’s instructions. Relative expression was calculated by the 2^–ΔΔCt method. Quantitative real-time PCR (qPCR) was performed using primer-probes for human MMP-9 (Hs00957562_m1, Thermo Fisher) per manufacturer’s instructions. qPCR was performed using an initial 10□min denaturation period at 95□°C followed by 40 cycles of 15□s at 95□°C and 1□min annealing and extension at 60□°C. Expression levels of MMP-9 were normalized to 18□S RNA (Thermo Fisher Scientific) while running simultaneously in each reaction. TaqMan is a probe-based assay that offers high specificity for target nucleic acids.

### Analysis of TNF-α, IL6 and IL1β by ELISA

Cell-culture supernatants were collected and stored at -80°C. Cell pellets were harvested in RIPA buffer containing protease inhibitors after the treatment period, clarified to remove debris, and analyzed using a human DuoSet TNF-α, IL6 and IL1β ELISA (R&D Systems) following the manufacturer’s protocol; absorbance was read at 450 nm on a BioTek Epoch. Data was normalized to a standard curve.

### β-Glucuronidase (GUSB) Activity

GUSB activity in supernatants was quantified using a commercial β-glucuronidase ELISA kit (Thermo Fisher Scientific) according to the manufacturer’s instructions; values were normalized to a standard curve.

### E2 Activity

Estrogen was measured in supernatants by human estradiol E2 ELISA Kit following the manufacturer’s protocol (Invitrogen); values were normalized to a standard curve.

### Statistical Analysis

All experiments were performed in triplicate and repeated independently at least three times. Data was analyzed using GraphPad Prism 9. Results are presented as mean ± SEM. Statistical significance was assessed using one-way ANOVA followed by Tukey’s post hoc test. A p-value < 0.05 was considered statistically significant.

## Results

### RSB11 suppresses MMP-9 expression across multiple cell models

To determine whether RSB11 in both its live and heat inactivated (HI) form modulates inflammatory protease activity, we measured MMP-9 expression in epithelial and non-epithelial cell types subjected to inflammatory injury. LPS stimulation (5 µg/mL, 4 h) or *E-coli* exposure (10^7^ CFU/mL) markedly increased MMP-9 mRNA in gut (Caco-2) **(Figure 1A)**, lung (HBE) **(Figure 1B)**, ovary (BG1) **(Figure 1C)**, bone (osteoblast MG-63) **(Figure 1D)**, kidney (A-498) **(Figure 1E)** and liver (HepG2) cells **(Figure 1F)**, confirming activation of inflammatory pathways in each model. Treatment with RSB11 at 3 mg/mL significantly suppressed MMP-9 expression across all cell lines tested. Importantly, the extent of suppression was indistinguishable between the live (RSB11-Life) and heat-inactivated (RSB11-HI) forms, with both preparations reducing LPS-induced MMP-9 to near-baseline levels. Both the live and HI formulations did not show any significant alterations in MMP-9 expression when treated in the absence of insults (LPS and E.coli) demonstrating their biosafety in these cell models. These findings clearly demonstrate that the ability of RSB11 to attenuate protease expression is not dependent on microbial viability.

**Figure 1.**
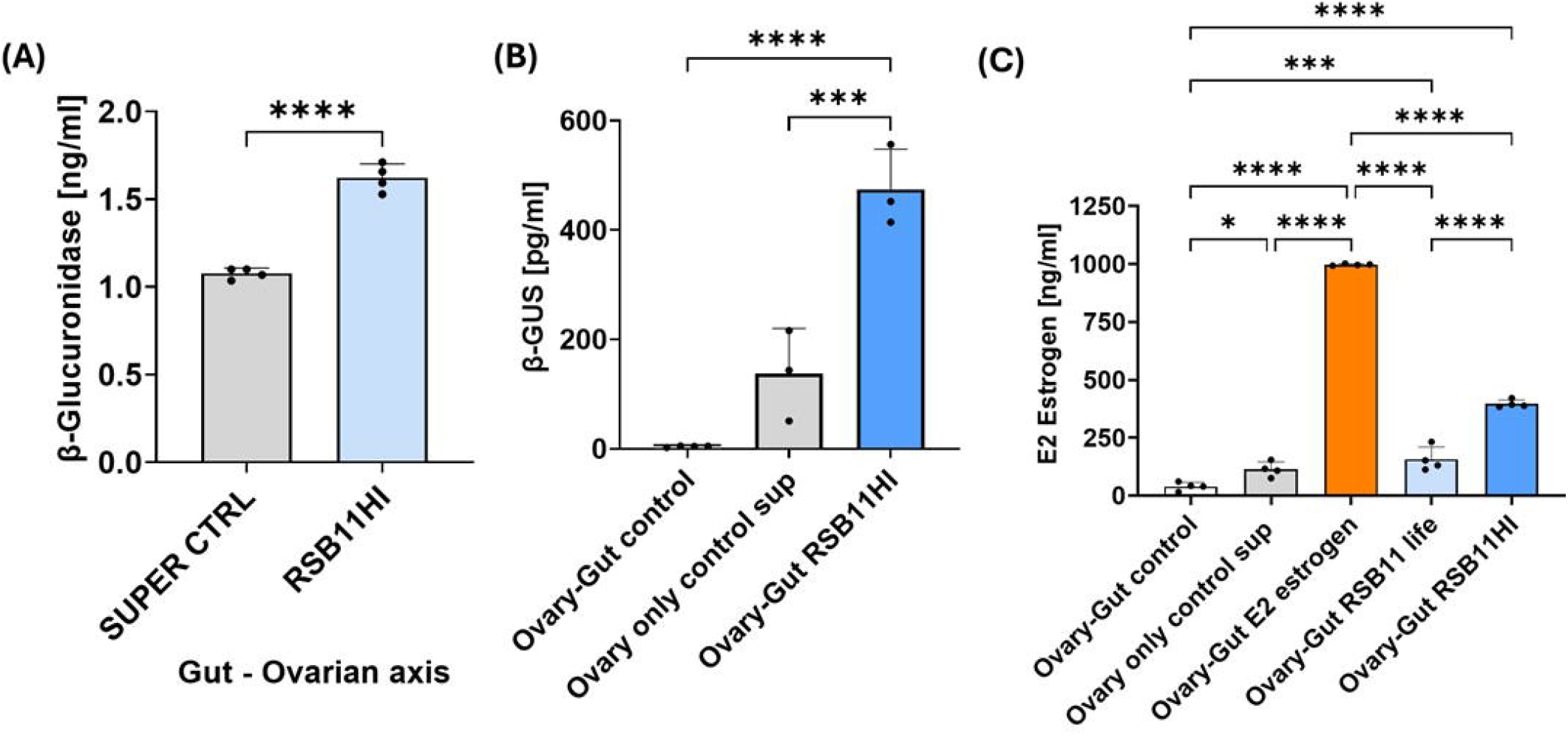
RSB11-Life and RSB11-HI suppress MMP-9 expression across multiple cell models. Quantitative PCR analysis of MMP-9 mRNA expression in. **(A)** Caco-2 (gut), **(B)** HBE (lung), **(C)** BG1 (ovary), **(D)** MG-63 (bone), **(E)** A-498 (kidney), and **(F)** HepG2 (liver) cells following stimulation with LPS (5 µg/mL, 4 h) or *E-coli* (10^7^ CFU/ml). Both live (RSB1 Life) and heat-inactivated (RSB11-HI) formulations (3 mg/mL) significantly suppressed MMP-9 induction to near-baseline levels, with no significant difference between the two forms. Neither RSB11-Life nor RSB11-HI altered basal MMP-9 expression in the absence of inflammatory challenge. Data was presented as mean ± SEM; n=3 independent experiments. p < 0.05, one-way ANOVA with Tukey’s post hoc test.

### RSB11 reduces TNF-α induction equivalently in live and heat-inactivated forms

Parallel analysis of cytokine production revealed that LPS and E.coli injury induced robust TNF-α, a pro-inflammatory cytokine, secretion in all cell types. Both RSB11-Life and RSB11-HI significantly reduced TNF-α levels, with reductions ranging from 40–70% depending on the cell type. Once again, no statistically significant difference was observed between live and inactivated preparations. The equivalence of effect was consistent across epithelial (Caco-2 **(Figure 2A)**, HBE **(Figure 2B)**, BG1**(Figure 2C)**) and non-epithelial (HepG2 **(Figure 2E )**, A-498 **(Figure 2H)**, MG63 osteoblasts **(Figure 2D)**) cell models in both the protein from cell pellet **(Figure 2A-E)** and supernatants **(Figure 2F-H)**, highlighting a broad anti-inflammatory profile retained after heat inactivation.

**Figure 2.**
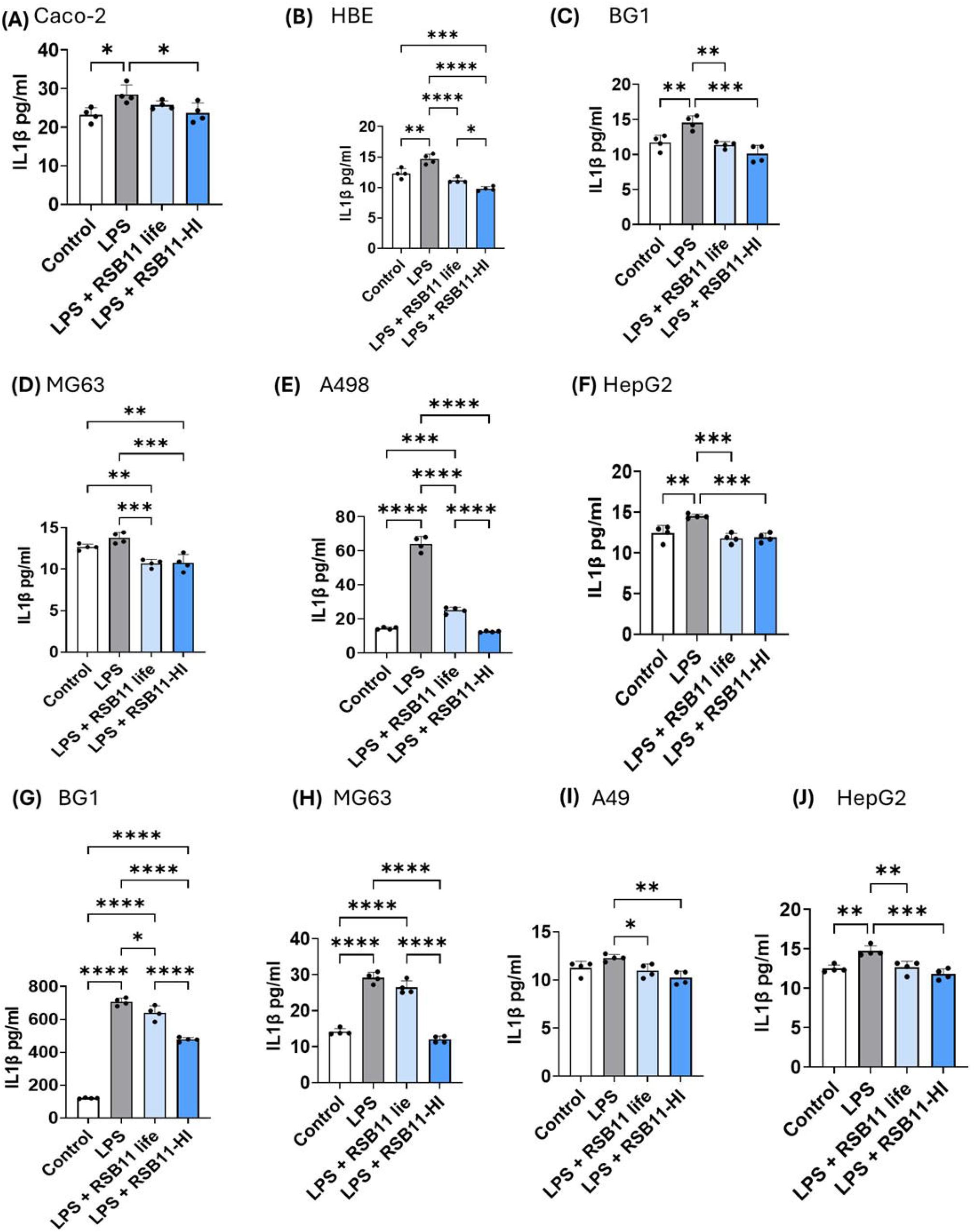
RSB11 life and RSB11-HI equivalently reduce TNF-α induction. ELISA measurement of TNF-α levels in cell lysates and supernatants from epithelial (A–C, F: Caco-2, HBE, BG1) and non-epithelial (**D–H**: HepG2, A-498, MG-63) cell models following LPS or *E-coli* challenge. Both RSB11 Life and RSB11-HI significantly reduced TNF-α secretion (40–70% reduction across models), with no significant difference between formulations. The equivalence of live and heat-inactivated preparations highlights the robust anti-inflammatory activity of RSB11 independent of microbial viability. Data shown as mean ± SEM; n=3 independent experiments. *p* < 0.05, one-way ANOVA with Tukey’s post hoc test.

### RSB11 and RSB11-HI attenuate IL-6 responses across multiple cell types

RSB11 life and its heat-inactivated form (RSB11-HI) demonstrated differential regulation of IL-6 across cell types. In Caco-2 and BG1 cells, neither RSB11 life nor RSB11-HI significantly suppressed IL-6 levels, intracellularly. However, in supernatants, only RSB11-HI significantly reduced IL-6 levels in BG1 cells **(Figure 3 A and C)**. In contrast, HBE cells exhibited a strong LPS-driven IL-6 increase, which was significantly reduced by both RSB11 life and RSB11-HI in cells **(Figure 3 B)**.

**Figure 3.**
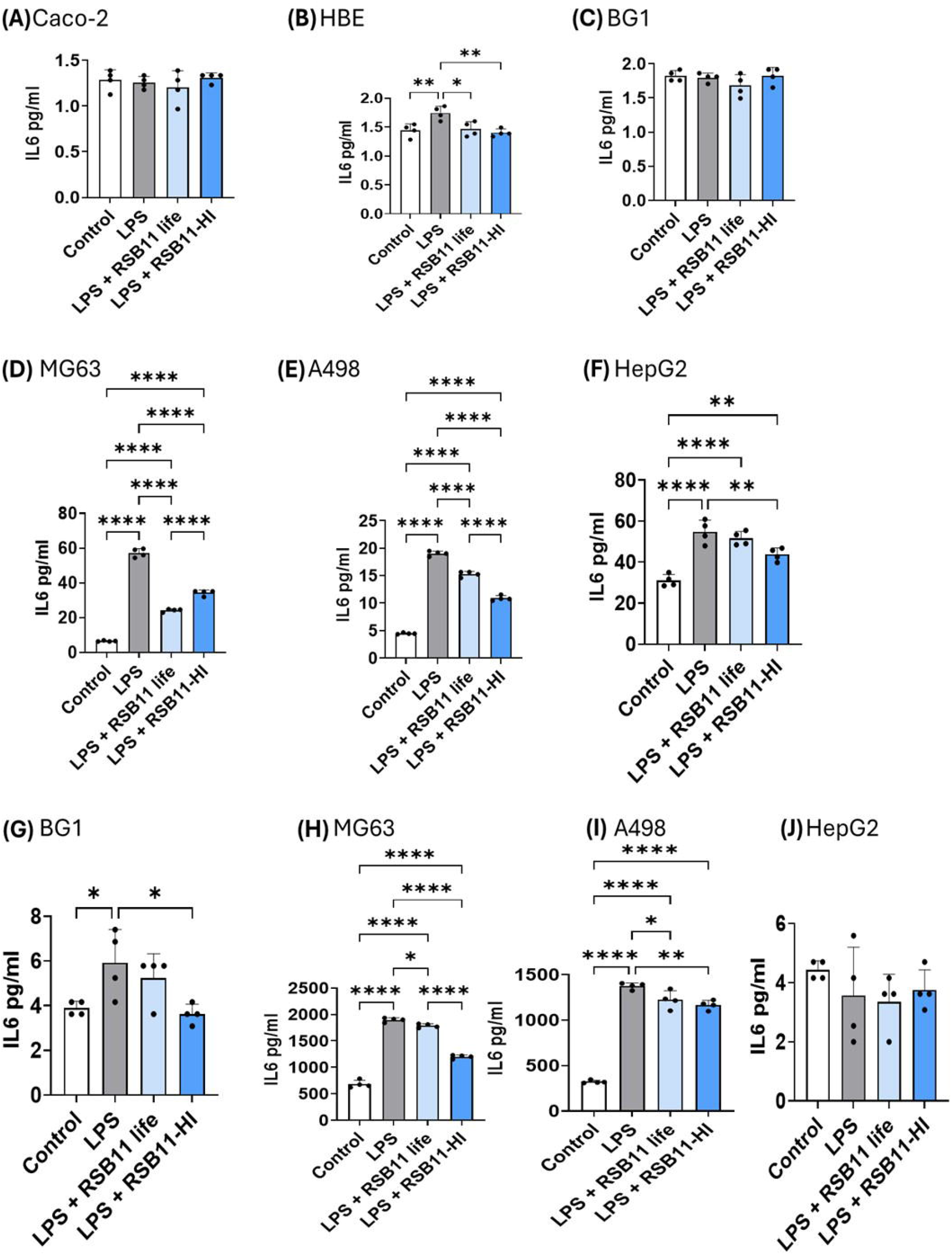
RSB11 life and RSB11-HI suppress IL-6 in (A-F) cell pellets and (G-J) supernatants. ELISA quantification of IL-6 levels in Caco-2, HBE, BG1, MG63, A498, and HepG2 cells. LPS stimulation significantly increased IL-6 expression across multiple cell types. RSB11 life and RSB11-HI significantly reduced IL-6 levels in HBE, MG63, A498, and HepG2 cells, both (A-F) intracellularly and in (G-J) supernatants, while little effect was observed in Caco-2 and BG1 cells. RSB11-HI produced stronger suppression than RSB11 life in MG63 supernatants. Data are mean ± SEM, n = 3–4, with p < 0.05, p < 0.01, p < 0.001.

The most pronounced effects were observed in MG63 and A498 cells. LPS markedly elevated IL-6 levels both intracellularly and in culture supernatants. Treatment with either RSB11 life or RSB11-HI significantly reduced IL-6 expression and secretion, with RSB11-HI consistently exerting stronger suppression in MG63 cells.

HepG2 cells also showed significant IL-6 reduction in cell pellets after treatment, though the supernatant levels were only modestly decreased. Collectively, these results indicate that both RSB11 life and RSB11-HI attenuate LPS-induced IL-6 responses in a cell-type-dependent manner, with robust suppression in osteoblast-like (MG63) and renal carcinoma (A498) cells **(Figure 3 D-J)**.

### RSB11 formulations mitigate IL-1β-driven inflammatory responses

Similar to IL-6, IL-1β, an inflammatory cytokine, responses varied among cell types. In Caco-2 cells, HBE cells and BG1 cells and supernatant, RSB11 life mitigated IL-1β levels however, RSB11-HI produced significantly higher suppression of intracellular IL-1β in all these cells **(Figure 4A-C, G)**. However, in MG63, A498, and HepG2 cells, both forms of RSB11 significantly reduced LPS-induced IL-1β expression both intracellular and secreted. A498 cells showed the strongest reduction, with IL-1β levels nearly normalized to control values following postbiotic treatment in cells, with the heat inactivated form showing the maximum efficacy in doing so **(Figure 4D-J)**. These findings indicate that both live and heat-inactivated RSB11 effectively blunt LPS-driven IL-1β responses across multiple cell types with the heat inactivated form being the most potent consistently.

**Figure 4.**
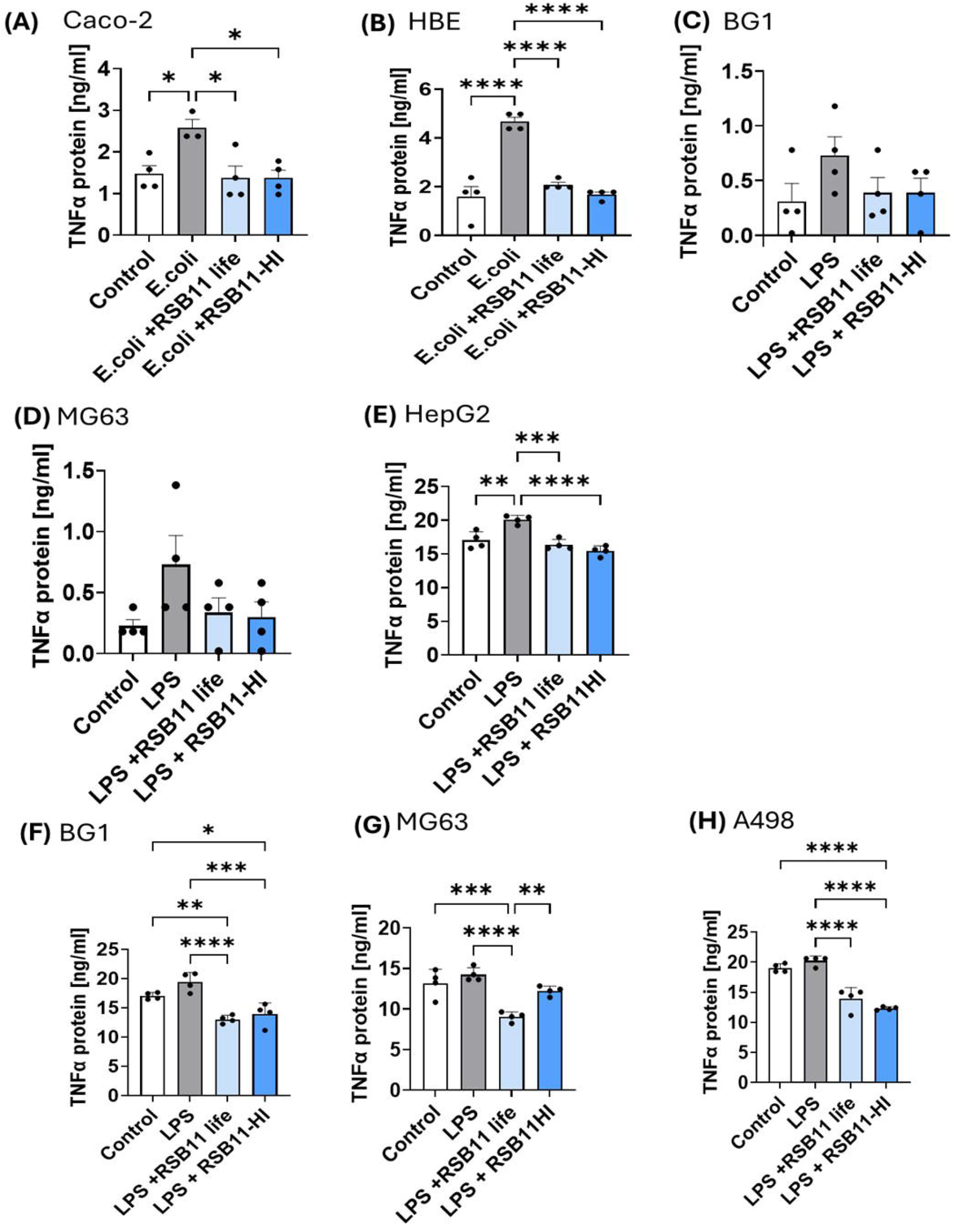
RSB11 life and RSB11-HI attenuate IL-1β in (A-F) cell pellets and (G-J) supernatants. ELISA measurement of IL-1β levels in Caco-2, HBE, BG1, MG63, A498, and HepG2 cells. LPS markedly increased IL-1β expression and secretion, which RSB11 life and RSB11-HI significantly reduced in HBE, MG63, A498, and HepG2 cells. A498 cells showed the strongest suppression, with IL-1β nearly restored to control levels. Minimal effects were observed in Caco-2 cells, while BG1 cells showed significant reductions only in secreted IL-1β. RSB11-HI frequently demonstrated slightly stronger suppression than RSB11 life. Data are mean ± SEM.

### RSB11-HI enhances β-glucuronidase activity and modulates estrogen levels along the gut–ovarian axis

To assess whether RSB11-HI influences metabolic pathways relevant to hormonal regulation, we quantified β-glucuronidase (β-GUS) activity and estrogen (E2) levels in gut–ovary co-culture models. Treatment with RSB11-HI significantly increased β-glucuronidase activity compared to control conditions **(Figure 5A)**. In gut supernatants, RSB11-HI nearly doubled β-GUS levels, while in the ovary–gut axis model, RSB11-HI also induced a robust increase in β-GUS compared to untreated controls **(Figure 5B)**.

**Figure 5.**
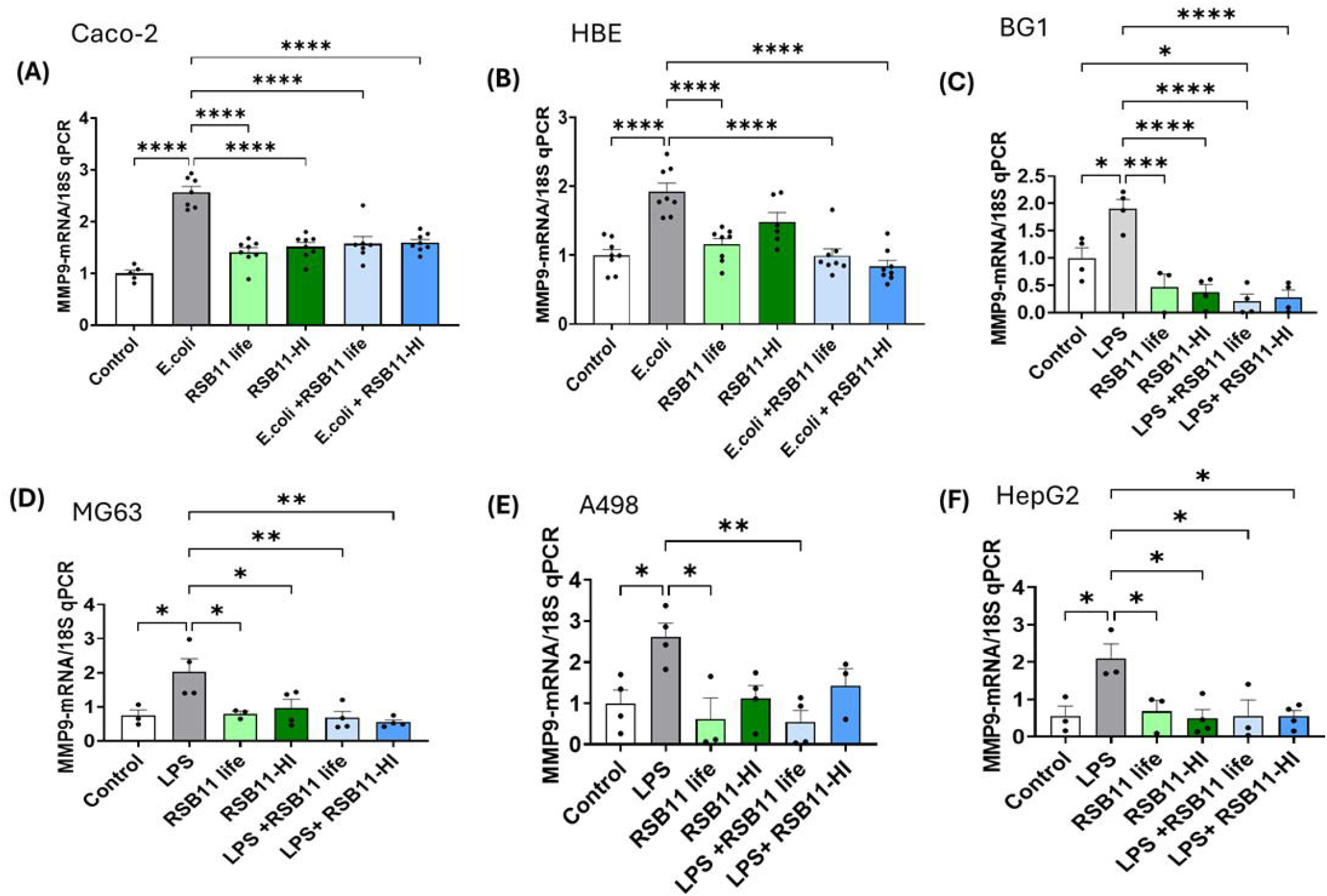
RSB11-HI enhances β-glucuronidase activity and estrogen production in gut–ovarian axis models. **(A)** β-glucuronidase (β-GUS) levels in gut supernatants were significantly increased following RSB11-HI treatment compared to supernatant controls. **(B)** In ovary–gut co-culture systems, RSB11-HI significantly upregulated β-GUS levels relative to untreated controls. **(C)** Estradiol (E2) levels were quantified across ovary– gut conditions. RSB11-HI markedly increased E2 levels compared to controls and RSB11life, while exogenous E2 supplementation produced the highest levels. Data are presented as mean ± SEM, n = 3–4, with p < 0.01 and **p < 0.0001 indicating statistical significance.

This enzymatic upregulation corresponded with a significant rise in estrogen production. In ovary–gut co-cultures, RSB11-HI treatment resulted in a ∼3-fold increase in E2 levels compared to control and RSB11-life treatments, indicating a stronger effect of the heat-inactivated form **(Figure 5C)**. Notably, exogenous estrogen supplementation validated the system, producing the highest E2 levels. These findings suggest that RSB11-HI not only suppresses inflammatory cytokines but also modulates metabolic-hormonal cross-talk along the gut– ovarian axis by enhancing β-glucuronidase activity and promoting estrogen production.

## Discussion

In this study, we applied the “gut-X axis” concept to outline the complex communication pathways between the gut and other distant organs (lung, liver, ovary, bone, kidney) and discussed our latest findings related to the effects of RSB11HI postbiotic on reducing inflammation and its implications in the gut-ovarian axis[33; 34; 35]. This bi-directional communication is mediated by factors like immune cells, hormones, and microbial metabolites, which can travel from the gut to these organs and influence their metabolic status, immune responses, and overall homeostasis[35; 36].

Previously we published the gut-lung axis where we tested an inhaled live biotherapeutic (LBP) using active Lactobacillus strains in *in vitro* and *in vivo* models of chronic lung diseases like BPD and COPD. The Lactobacillus-based LBP is effective in improving lung structure and function, mitigating neutrophil influx, and reducing a broad swath of pro-inflammatory markers in these models of chronic pulmonary disease via the MMP-9/PGP (matrix metalloproteinase/proline-glycine-proline) pathway[11; 37].

In this in vitro study we extend our research to other organs and compare the live biotherapeutic (RSB11 life) to its heat-inactivated form (RSB11-HI), a Resbiotic Nutrition Inc. proprietary form[22; 32]. Understanding these gut-X-axes opens new avenues for treatment, such as using postbiotics or dietary interventions to target the gut microbiota and improve conditions in other organs[34; 35; 38]. The therapeutic potential of RSB11-HI may have forthcoming clinical relevance in various diseases, including metabolic disorders, mental health conditions, and autoimmune diseases by potentially reducing systemic inflammation.

Current data provides compelling evidence that the postbiotic RSB11 retains potent immunomodulatory and metabolic functions even after heat inactivation. Both live RSB11 and its heat-inactivated counterpart (RSB11-HI) significantly attenuated pro-inflammatory cytokines, including TNF-α, MMP-9, IL-6, and IL-1β, across diverse cell types representing gut, lung, ovary, bone, kidney, and liver systems. Importantly, RSB11-HI was not only equivalent to the live form in suppressing these inflammatory mediators but, in some contexts, demonstrated superior efficacy. Beyond immunomodulation, RSB11-HI uniquely enhanced β-glucuronidase activity and estrogen production along the gut–ovarian axis, expanding its functional profile to include host– microbiome hormonal cross-talk. These findings reframe the paradigm of probiotic therapeutics by showing that microbial viability is not essential for therapeutic benefit, and that structural or metabolite-retained components of postbiotics may provide a safer, more versatile alternative. This study employed a heat-treated preparation of RSB11 produced under controlled thermal conditions to ensure consistency and stability. The comparable biological efficacy observed between the live and heat-treated forms highlights the robustness and reproducibility of the RSB11 formulation.

The ability of RSB11-HI to modulate β-glucuronidase and estradiol levels expands its relevance beyond inflammation. β-Glucuronidase is essential for estrogen deconjugation and recirculation; its dysregulation has been linked to estrogen deficiency and reproductive dysfunction [6; 39]. By enhancing this pathway, RSB11-HI may restore estrogenic tone in disorders such as PCOS, menopause-associated bone loss, and estrogen-deficiency mood disorders. The integration of endocrine modulation with anti-inflammatory activity differentiates RSB11-HI from conventional postbiotics, suggesting unique value in women’s health and broader metabolic diseases.

Our cytokine analyses confirmed that RSB11-HI exerts broad anti-inflammatory effects[34; 40]. IL-6 and IL-1β, both pivotal drivers of acute inflammation and chronic disease, were robustly induced by LPS in epithelial, hepatic, renal, and osteoblast-like cells. Treatment with RSB11-HI consistently reduced both intracellular and secreted cytokine levels, with the strongest suppression observed in MG63 and A498 models. These findings complement the previously demonstrated inhibition of TNF-α and MMP-9, together establishing that RSB11-HI targets multiple inflammatory nodes[34; 40]. The ability to concurrently suppress proteases, cytokines, and upstream drivers of inflammation positions RSB11-HI as a broad-spectrum immunomodulator with potential utility in conditions such as inflammatory bowel disease, arthritis, pulmonary fibrosis, and cancer, where these mediators are central to pathology.

A major barrier to probiotic therapy is the requirement for microbial viability [41; 42]. Live probiotics are prone to loss of efficacy during storage and distribution, necessitate strict cold-chain logistics, and can pose safety concerns in vulnerable or immuno-compromised populations[14; 43]. Our findings that RSB11-HI is fully active and, in some cases, more potent than its live form demonstrate that heat inactivation does not compromise function but instead provides distinct advantages. By eliminating the dependence on microbial survival, RSB11-HI allows for stable formulations with extended shelf-life, reduced manufacturing costs, and compatibility with diverse delivery platforms (capsules, powders, beverages, or topicals). Furthermore, safety is inherently improved, removing concerns of uncontrolled replication, horizontal gene transfer, or sepsis in high-risk patients[44; 45]. Thus, RSB11-HI represents a paradigm shift from “live-dependent” probiotics to next-generation postbiotics optimized for safety, stability, and scalability.

One of the most striking findings of this study is the ability of RSB11-HI to modulate β-glucuronidase activity and estrogen metabolism along the gut–ovarian axis. β-Glucuronidase is a key microbial enzyme involved in the enterohepatic recirculation of estrogens[46]. By enhancing β-glucuronidase activity, RSB11-HI increased estrogen bioavailability in ovary–gut co-culture models, leading to significantly elevated estradiol levels compared to control or live RSB11 treatment. This dual action, dampening inflammation while restoring estrogen signaling, suggests that RSB11-HI may influence a broader network of host–microbiome interactions that extend beyond classical immune pathways. Dysregulation of estrogen metabolism is implicated in reproductive health disorders, osteoporosis, cardiovascular disease, and hormone-driven cancers[47]. Thus, the ability of RSB11-HI to modulate both immune and hormonal pathways positions it uniquely as a therapeutic candidate at the intersection of inflammation and metabolism

From a translational perspective, the equivalence and occasional superiority of RSB11-HI relative to the live form opens multiple therapeutic avenues[48]. The combined suppression of TNF-α, IL-6, and IL-1β mirrors the targets of several high-value biologics currently used in clinical practice, such as anti-TNF and anti-IL-6 therapies. Moreover, the capacity to enhance β-glucuronidase activity and estrogen availability introduces opportunities for application in women’s health, including conditions such as polycystic ovary syndrome (PCOS), infertility, menopause-related osteoporosis, and estrogen-linked mood disorders. The integration of immune dampening and hormonal regulation is especially relevant to diseases where chronic inflammation and endocrine imbalance converge, such as metabolic syndrome, autoimmune conditions, and hormone-driven cancers.

From a translational perspective, the equivalence of live and HI forms on inflammation, combined with the unique hormonal modulation of RSB11-HI, positions it for dual-action indications. Unlike biologics (e.g., anti-TNF, anti-IL-6 antibodies), which are costly and immunosuppressive, RSB11-HI represents a safe, orally deliverable, and cost-effective intervention[49; 50]. Its dual targeting of inflammation and estrogen balance may be especially valuable in female-predominant diseases where endocrine dysfunction exacerbates immune activation (e.g., autoimmune thyroiditis, endometriosis, rheumatoid arthritis).

This study, while comprehensive across multiple epithelial and non-epithelial in vitro models, remains limited to cell culture systems. Although these models capture key aspects of inflammatory and hormonal signaling, they cannot fully recapitulate the systemic complexity of host–microbiome interactions *in vivo*. Future validation in animal models and clinical cohorts will be vital to confirm the translational relevance of RSB11-HI, particularly in the context of enterohepatic circulation and endocrine cross-talk. Moreover, expanding analyses to include long-term dosing paradigms, microbiome compositional shifts, and tissue-specific hormone receptor activation could further strengthen mechanistic insight. Consistent with these findings, supplementation with purified bacterial cell wall components such as teichoic acid, peptidoglycan, polysaccharide, and S-layer, each independently attenuated MMP-9 expression in E. coli-challenged bronchial epithelial cells[37]. This suggests that structural components of Lactobacillus strains contribute significantly to the anti-inflammatory activity of RSB11-HI. Such results further support the hypothesis that postbiotic efficacy arises not solely from secreted metabolites but also from stable, cell-associated molecular patterns capable of modulating host immune signaling. The strong and consistent effects observed across multiple independent cell models provide a robust foundation for advancing RSB11-HI into preclinical development, supporting its potential as a dual-action postbiotic with both anti-inflammatory and endocrine-modulatory benefits.

In summary, this study shows that heat-inactivated RSB11 is a potent and versatile postbiotic that suppresses key inflammatory mediators (MMP-9, TNF-α, IL-6, IL-1β) while enhancing β-glucuronidase activity and estrogen bioavailability. A multifaceted therapeutic modality of immune suppression and hormonal modulation underscore the breadth of its therapeutic potential across gastrointestinal, pulmonary, reproductive, hepatic, and metabolic diseases. By combining the functional efficacy of live probiotics with the superior safety, scalability, and translational feasibility of inactivated preparations, RSB11-HI emerges as a next-generation microbial therapeutic with broad clinical relevance.

## Data Availability Statement

Conflicts of Interest: Part of the research described in this manuscript is patented under “Inhaled Respiratory Probiotics for Lung Diseases of Infancy, Childhood and Adulthood” US 11,141,443 B2 held under the University of Alabama at Birmingham Research Foundation (C.V.L, A.G., N.A. are inventors). The remaining authors declare no competing interests.

### Abbreviations

BPD: Bronchopulmonary Dysplasia
HI: heat inactivated
COPD: Chronic obstructive pulmonary disease
TNF-α: Tumor necrosis factor-α
IL-6: Interleukin-6
IL-1β: Interleukin-1β
MMP-9: matrix metalloproteinase-9

## Author contributions

Conceptualization: T.N. and C.V.L. Experimentation: T.N., T.M., A.M., Y.Y., I.A., and R.S. Analysis and interpretation: T.N., A.A., T.M., K.S., N.A. and C.V.L. Drafting and editing manuscript: T.N., A.A., T.M., K.S, N.A. and C.V.L. Funding acquisition and research supervision: T.N., N.A. and C.V.L. All authors have approved the final version of the manuscript as submitted.

